# CoverageMaster: comprehensive CNV detection and visualization from NGS short reads for genetic medicine applications

**DOI:** 10.1101/2021.07.21.453195

**Authors:** Melivoia Rapti, Jenny Meylan, Emmanuele Ranza, Stylianos E. Antonarakis, Federico A. Santoni

## Abstract

CoverageMaster (CoM) is a Copy Number Variation (CNV) calling algorithm based on depth-of-coverage maps designed to detect CNVs of any size in exome (WES) and genome (WGS) data. The core of the algorithm is the compression of sequencing coverage data in a multiscale Wavelet space and the analysis through an iterative Hidden Markov Model (HMM). CoM processes WES and WGS data at nucleotide scale resolution and accurately detect and visualize full size range CNVs, including single or partial exon deletions and duplications. The results obtained with this approach support the possibility for coverage-based CNV callers to replace probe-based methods such array CGH and MLPA in the near future.

## Introduction

Copy number variation (CNV) is the most frequent structural alteration in the human genome. Aberrant numbers of copies of specific genes, exons or, in general, genomic regions are known to be implicated in pathogenic conditions such as Mendelian diseases and cancer ^1–4^. Hence, identification of these deletion and amplification events is a primary purpose in medical genetics research. In clinical diagnostics, the identification of rare, potentially causative CNVs in a patient with a suspected genetic disorder is a long-sought objective. However, the discovery of such mutations that can vary in size and copy number is a challenging task. Currently the most commonly used methodologies to detect clinically relevant CNVs rely on microarray based technologies. Array comparative genomic hybridization (array CGH) offers an efficient method to detect CNVs and micro-CNVs (5kb < size < 10Mb) in the whole genome, but its resolution is not covering the lower size spectrum. Multiplex ligation-dependent probe amplification (MLPA) is the current golden standard to detect exon-sized CNVs but this technology can cover few exons per assay (low throughput) and its application is limited to a small number of genes ^5^.

In recent years, the development of next generation sequencing (NGS) technologies of short reads has provided a standardized way for accurate coding variant analyses through whole genome sequencing (WGS) and whole exome sequencing (WES). Remarkably, this technology provides the coverage per nucleotide of clinically relevant regions of the genome. Whereas WGS allows for a more comprehensive overview of the entire genome with uniform coverage^6^, the related sequencing costs and the computational infrastructures needed to process the raw data are still limiting its broad application in clinical practice^7^. On the other hand, WES is computationally less demanding and has reached such a high sensitivity and specificity in variant calling to eventually become a clinical standard. Currently, WES is widely used for diagnostic purposes in many medical genetics laboratories throughout the world.

A wide range of detection algorithms have been developed to call CNVs from WGS and WES data. Read-depth based methods are considered the most effective for accurate copy number prediction and exploit the fact that NGS generates raw data in the format of short reads^11^. These reads are mapped to a reference sequence and the coverage in a genomic region is calculated by counting the number of reads that align to this region. Depth-of-coverage (DoC) is then assumed to be proportional to the copy number of that region. In principle, DoC is sufficient for the detection of all clinically relevant CNVs, irrespectively of size and copy number, promoting WGS and WES as a robust and more inclusive alternative to complementary laboratory approaches such as array CGH or MLPA.

Nevertheless, WES has technical issues that result in the generation of noisy data. First, the lack of continuity of the target regions and, second, the biases due to hybridization and sequencing processes complicate the procedure to standardize CNV detection^11^. As a result, current WES based detection methods suffer from limited resolution, high false positives and false negatives calls.

Here, we introduce CoverageMaster (CoM), a CNV calling algorithm based on depth-of-coverage maps from aligned short sequence reads from WES or WGS. CNVs are inferred with Hidden Markov Models at multiscale nucleotide-like levels in the Wavelet reduced space, in comparison to existing methods that utilize fixed length windows or exon averages. This approach is designed to optimize the search for CNVs of different sizes. Working at nucleotide resolution, CoM provides the graphical representation of the predicted CNV in all genes of interest, and, optionally, a wig formatted file compatible with UCSC Genome Browser for detailed visualization of the normalized coverage on the target genes or regions in the genomic space. We propose CoverageMaster as a potential first-line diagnostic tool in research and clinical applications.

## Results

CoverageMaster utilizes the representation of coverage signal ratio (case over control) in the reduced Wavelet approximation space to perform a multiscale analysis of aberrant coverage profiles, potentially underlying causative CNVs, at nucleotide resolution (Figure 1, see Methods). This approach is meant to explore a broad spectrum of CNV sizes and in particular deletions or duplications of < 5kb. At this scale, the experimental noise is caused on one hand by the particular technology used for sequencing and, for WES, DNA selection by hybridization. On the other hand, batch specific coverage distortions may occurs. Intuitively, the smaller the CNV the higher the chance that the call is a false positive. To overcome this problem, CoverageMaster exploits the fact that, as all other genomic variations, clinically relevant CNVs are rare (MAF<0.01%). Thus, it is reasonable to assume that such CNVs cannot be present in two or more independent unrelated individuals of the same batch. Following this basic principle, CoM utilizes a reference with the average coverage and standard deviation of 15-20 samples processed with the same technology (hybridization kit, reagents, sequencer). The reference provides the standard deviation per nucleotide from the expected coverage where coverage spikes are produced by reproducible experimental noise and/or recurrent CNVs. Eventually, matching CNVs in the test sample are then considered as frequent or false positives and finally discarded. Moreover, CoM pairwise compares the sample case with independent samples, used as controls, from the same batch. Spikes present in the test signal coverage and in one control sample are averaged out in the coverage ratio and consequently discarded.

**Figure 1.**
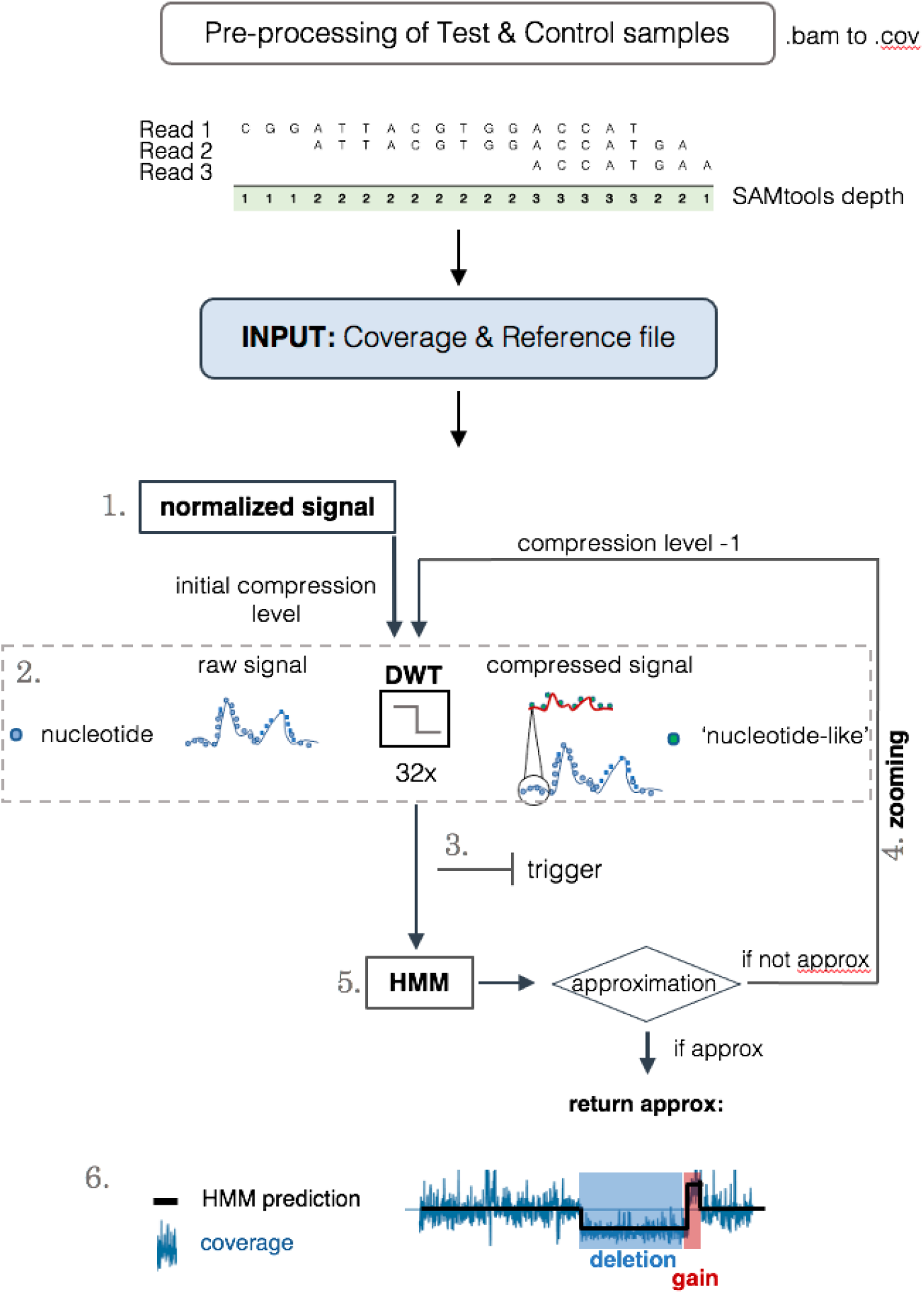
CoverageMaster workflow. CoM is based on depth-of-coverage maps from aligned short sequence reads from WES or WGS. The normalized values of the depth-of-coverage for each nucleotide position are calculated (Step 1). The ratio of the test to control coverage signal is compressed at a specified initial scale *l* by default in the nucleotide-like space using the Discrete Wavelet Transform (DWT) (Step 2). For the compressed signal, an indicator detects the potential nondiploid nucleotide-like positions (Step 3). HMM is used to segment the compressed signal into regions of similar copy numbers and assign CNV states (Step 4). If no putative CNVs are identified, the process is repeated at scale *l* – 1 via “zooming” (Step 5).

In order to prove its efficiency, we tested CoverageMaster in various contexts of WES. All samples processed here were hybridized with Twist Core Exome + RefSeq Spike and sequenced with Illumina HSeq4000 or Novaseq.

To demonstrate the performance of CoM in standard clinical analyses, we analyzed 12 clinical samples and compared CoM CNVs calls to standard array CGH calls (see Methods). In order to provide a point of reference, we also included ExomeDepth (ED), a performant DoC-based CNV caller ^10^. In Figure 2a, the cumulative true positive values for CNVs detected by CoM and ED are reported. CoM calls coincide with almost all of the array CGH calls for each sample with the exception of some frequent benign variants discarded by CoM because they were present in most controls. In fact, when searching for CNVs with MAF<1%, CoM identifies all CNVs detected by array CGH, in contrast to ED that detect 80% of them (Figure 2b). This result demonstrates that CoM may replace array CGH in clinical diagnostic settings.

**Figure 2.**
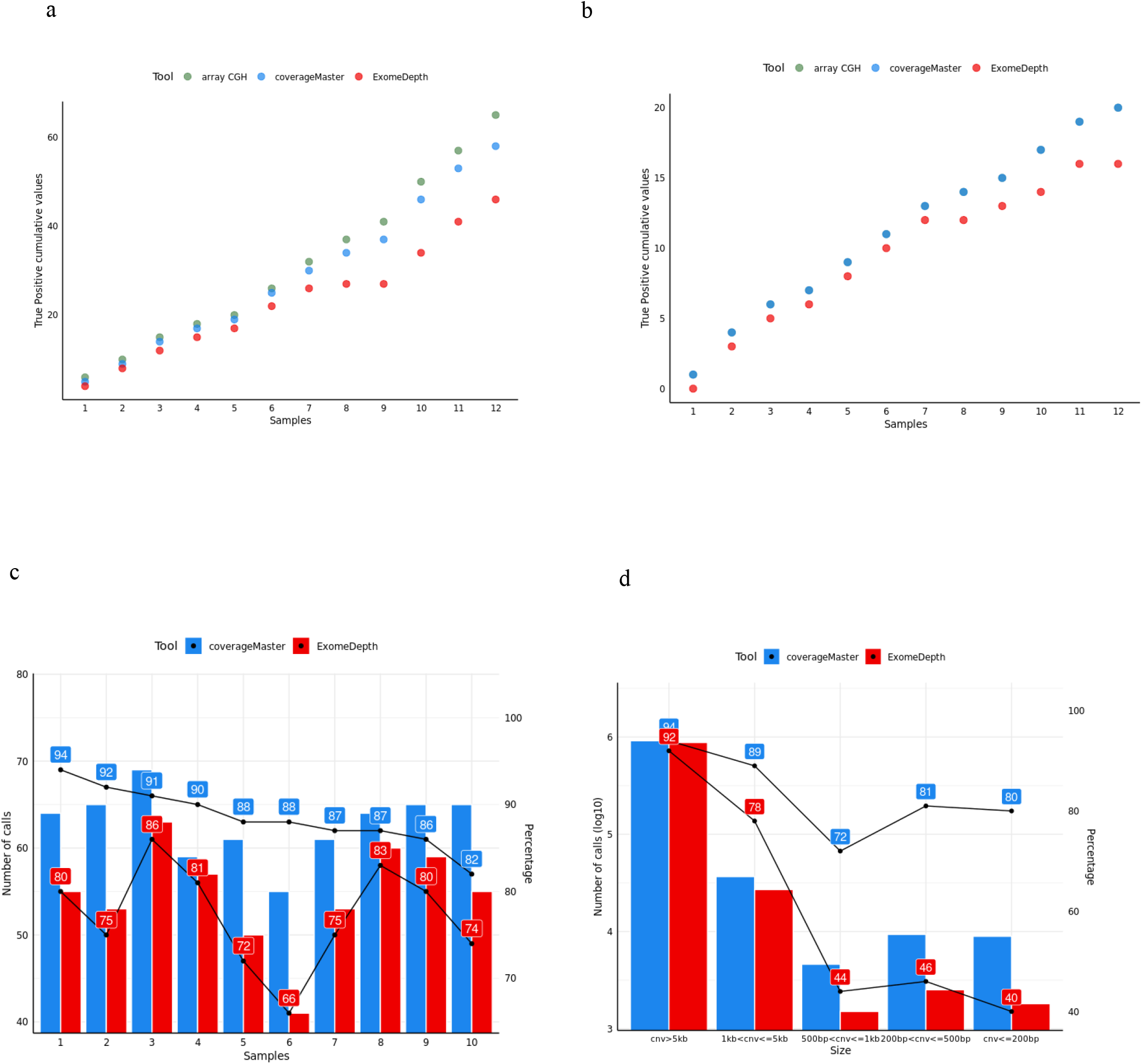
Array CGH comparison on clinical samples and CNV detection in simulated data. Cumulative plots of number of calls (y-axis) detected by CoM (blue) and ED (red) and the number of CNVs found by array CGH (green) in 12 samples: a) all calls are considered; b) only the rare CNVs (MAF<1%) are included, green and blue dots are overlapping; c) number and fraction of true calls detected by CoM (blue) and ED (red) in 10 samples where CNVs of various size were randomly introduced in exonic regions; d) number and fraction of true calls of detected CNVs stratified by size for CoM (blue) and ED (red).

To investigate the performance of CoM on small CNVs, given the lack of a standard test set for CNV detection of this size, we created a dataset of simulated WES data from 10 individuals. Around 700 heterozygous duplications and deletions of 200, 500, 1000 and 5000 base pairs were randomly introduced in the exonic regions of each sample (see Methods) and analyzed by CoM and ED. The results show that CoM achieves a total sensitivity of 90% as compared to 77% obtained by ED (Figure 2c) and an average precision of 41% for CoM versus 37% obtained by ED (an average of ~270 calls out of 18000 genes for both programs, precision is calculated considering half of the calls as false positives given there is no validation available). Most importantly, CoM can accurately detect deletions and duplications of (< 200bp) with sensitivity above 80%, when the corresponding percentage for ED is 40%. Notably, the multiscaling approach allows CoM to keep a similar performance along all size classes while ED rapidly decreases its performance with size reduction (Figure 2d).

CoM has been mainly conceived as a diagnostic support tool for clinical genetics analysis. To provide a perspective of the broad capabilities of the algorithm, we report three examples of solved clinical cases.

Patient 1 is a 20 years old male presenting with a congenital obesity of class III. WES analysis did not provide any suitable SNV or INDEL candidate variant on a panel of 48 genes for monogenic obesity (https://www.medigenome.ch/en/gene-panels/). On the same panel, CoM identified a heterozygous deletion of ~200kb in SH2B1 (Figure 3a). This deletion is known as the chromosome 16p11.2 deletion syndrome [OMIM 613444] and is a common cause of congenital obesity. The CNV was subsequently confirmed by array CGH.

**Figure 3.**
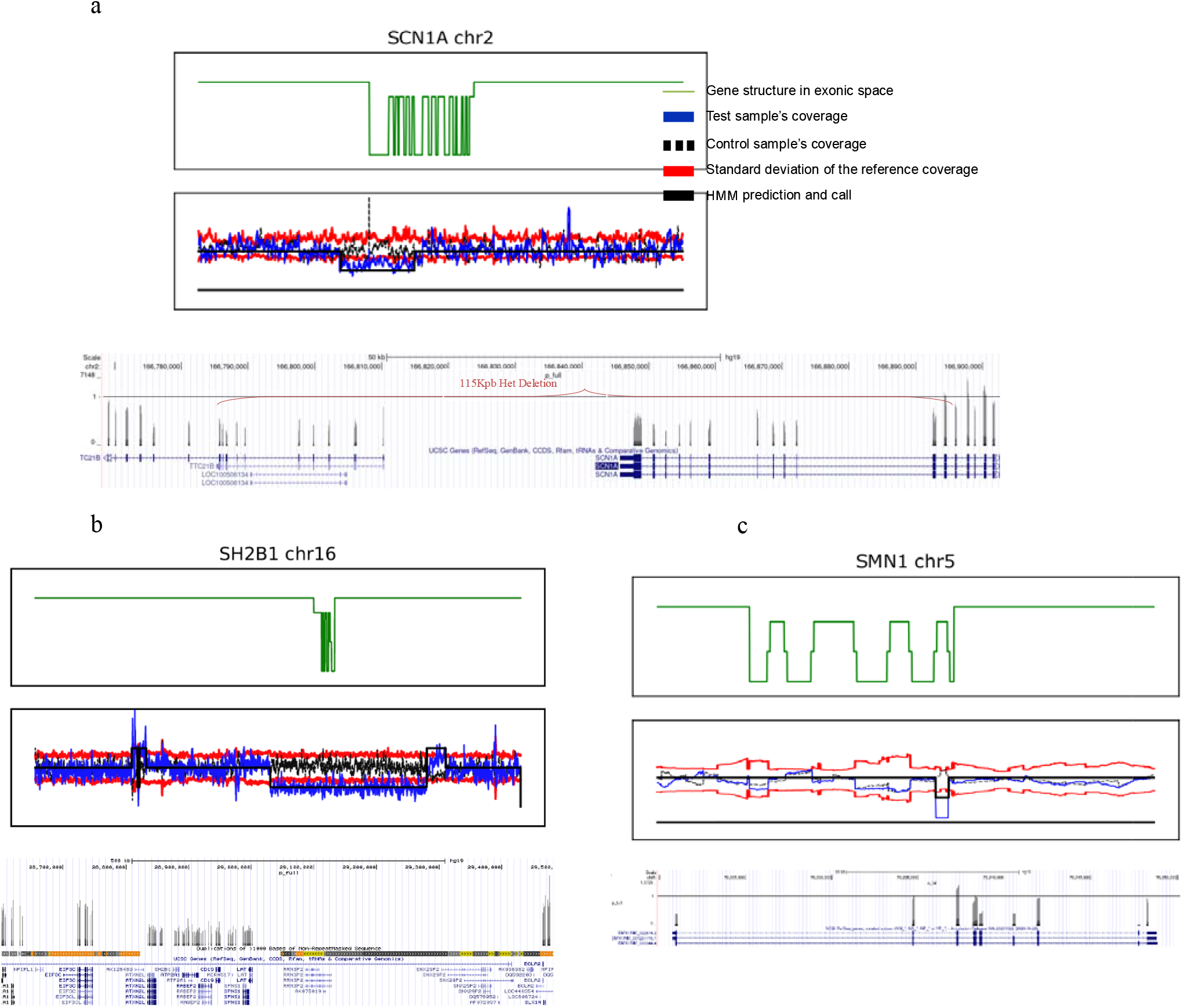
Examples of clinically relevant CNV identified by CoM. a) In green, the collapsed exon structure of the gene of interest, up or down blocks representing one exon. Coverage profiles in exon space of test sample, control and reference (color code in the legend) are represented in the second plot. For patient 1 (see text), the partial heterozygous deletion of 115kb in SCN1A is clearly visible in the exonic and genomic spaces. b) Heterozygous deletion encompassing SH2B1 detected in patient 2 (see text). Being in a region of repeated sequences (lower plot in the genomic space), the size of this deletion is not clearly ascertainable. c) Homozygous deletion of exon 7 in SMN1 in patient 3 clearly visible in the exonic and genomic spaces. It is worth noting that, in the genomic space, the coverage profile seems to show two other exons with a drop in coverage. The control, dashed line in the plot above, shows the same profile indicating a fluctuation of the coverage in this region, likely independent from the number of copies.

Patient 2 is an 8 years old female child, diagnosed with drug-resistant epilepsy with febrile seizures. WES analysis did not provide any candidate variant on a panel of 478 genes related to epilepsy (Epilepsy MDG-1204.01, https://www.medigenome.ch/en/gene-panels/). CoM reported a heterozygous deletion of ~120kpb partially overlapping the last 10 exons of SCN1A (Figure 3b). The sodium channel 1A is associated with generalized epilepsy with febrile seizures, Type 2 [OMIM 604403]. Deletions in this gene are known to cause seizure disorders, ranging from early-onset isolated febrile seizures to generalized epilepsy^12^.

Patient 3 is a 3 years old female child with a suspicion of spinal muscular atrophy. WES analysis and array CGH were negative but CoM identified a full exon 7 homozygous deletion of 112bp (Figure 4c). This deletion, which was confirmed by MLPA and not detected by ED, is the most frequent CNV related to SMN1 induced muscular atrophy^13^ [SMA OMIM 253400]; this deletion was therefore considered as the pathogenic cause of the phenotype of the patient by the clinicians.

## Discussion

CoverageMaster is a NGS coverage based CNV calling algorithm designed to work at nucleotide resolution with WES or WGS data. The capacity to analyse a given coverage signal in different scale sizes, combined with the nowadays availability of numerous controls in standard clinical batches, enables the detection of multi-sized clinically relevant deletions or duplications and in particular the detection of the so far elusive small CNVs of < 5kb. We have proven the effectiveness of CoverageMaster in comparison to another excellent and broadly used *in silico* CNV caller, ExomeDepth. Performance wise, CoM takes 1h20min to process a panel of 4758 clinically relevant genes from OMIM and the Clinical Genomic Database (https://research.nhgri.nih.gov/CGD/) on a 16 cores machine with 32Gbyte of RAM. CoM demonstrated to be superior to ED in the detection of rare and small CNVs in simulated and clinical data and to be a valid and inexpensive alternative to MLPA and array CGH in clinical settings.

## Materials and Methods

The analyses reported in this study were performed on DNA processed by whole exome sequencing at the Health 2030 Genome Center (https://www.health2030genome.ch/) or Medigenome (www.medigenome.ch) using Twist Human Core Exome Kit (TWIST Biosciences, San Francisco, CA, USA); sequencing was performed on Illumina HiSeq4000 or Novaseq platforms. Array CGH and MLPA were performed in GeneSupport using Agilent SurePrint G3 Human 4×180K and SALSA MLPA Probemix P021 SMA (MRC Holland), respectively.

### Preprocessing and transformation of exome data

CoverageMaster (CoM) uses DoC maps from aligned short sequence reads to estimate CNV events. To acquire the sequence reads, the mapping is done with the standard pipeline for whole exome or whole genome sequencing data based on GATK^14^, and the coverage at each nucleotide of the region of interest (ROI) is calculated and stored in tab separated COV files (format: chr nucleotide_position coverage) using samtools (samtools depth)^15^. Coverage files of a test/target plus one or more controls plus one reference coverage serve as input for the algorithm. The assumption is that control coverages are DoC maps of copy number neutral cases (diploid) or carrier of frequent CNVs in the ROI of interest. The reference set consists of a batch of coverage files from samples processed with the same technology (i.e. hybridization kit, reagents for library prep and sequencer) used to generate case and controls. First, the coverage per nucleotide per sample is normalized by the respective total number of reads. Then, mean and standard deviation of the normalized coverage values are computed over all the samples for each nucleotide.

### Wavelet transform

In a genomic region of *N* nucleotides, the coverage of test case and control can be represented as the discrete signals *s*(*n*) and *c*(*n*), respectively, where *n* is the nucleotide number corresponding to the genomic or exonic position in the exon space (the space where covered regions (i.e. exons) are “ligated” together). In the ideal case, the coverage ratio 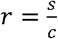 is a non-periodic square waveform with up and down steps in correspondence of increased or decreased copy number, respectively. In order to diminish the noise induced by fast variations of the signal and, at the same time, to reduce the computational burden, the coverage ratio is compressed in the nucleotide-like space using the Discrete Wavelet Transform (DWT) equipped with the Haar basis. At scale *l,* the approximation and detail coefficients are *r_l_, d_l_, d*_*l*-1_,…, *d*_0_ = *DWT_l_*(*r*). The *M* = *N* · 2^-*l*^ approximation coefficients *r_l_* are normalized to the median of the original signal and used for CNV analysis.

### Multiscale CNV detection

The probability *b_j_*(*o_m_*) of each *m* nucleotide-like positions of the sequence of approximation coefficients *r_l_* = *o*_1_*o*_2_… *o_k_o*_*k*+1_…*o_M_* to be in a normal (i.e. diploid), duplicated or deleted state 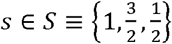 is defined at any scale *l* as a random variable with Gaussian distribution of mean *s* and standard deviation *σ*(*R_l_*), where *R_l_*(*m*) is the sequence of approximation coefficients of the reference coverage in the *m*-coordinates of the *l*-scaled nucleotide-like space.

At scale *l,* the indicator function (“trigger”) *T* = *argmax_s_*(*b_s_*(*r_l_*)) ≠ 1 identifies the locations of non diploid nucleotide-like positions and masks the rest of the signal. If no location is identified, the algorithm discards this region and processes the next one.

Once the putative CNVs are identified, the Viterbi algorithm is then used to identify the most likely copy number state sequence *Q* = *q*_1_*q*_2_…*q_k_q*_*k*+1_…*q_M_* of the compressed genomic region, based on the corresponding sequence of observations 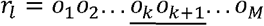. Masked observations 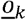 have a fixed diploid state *q_k_*=1.

More formally, if *v_t_*(*j*) represents the Viterbi probability that the underlying HMM is in copy number state *j* after seeing the first *m* observations and passing through the most probable state sequence *q*_1_*q*_2_… *q*_*m*-1_, it can be shown that

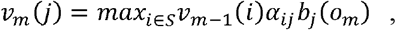

where *v*_*m*-1_(*i*) is the previous Viterbi path probability from the previous nucleotide, *a_ij_* is the transition probability (here set to *a_ij_* = 5 · 10^-6^ which is the probability of finding a duplication or a deletion in the human genome, calculated as the mean of the inclusive and stringent number of CNVs per nucleotide from ^16^) and *b_j_*(*o_m_*) is the observation probability given the state *j* as defined above.

If no putative CNV is detected at this stage, the algorithm performs a multiscale analysis by repeating the HMM phase with the masked signal transformed at scale *l* – 1. Again, in absence of CNVs, the algorithm keeps decrementing *l* down to, if necessary, *l* = 0 (no compression). This is computationally possible because only the relevant unmasked regions are actually inspected. Otherwise, eventual putative CNVs are saved and the algorithm proceeds to the next region.

### Iteration over controls

In case more control coverages are provided, eventual putative CNVs and relative masks are stored in a temporary buffer. Following the assumption that a rare causative CNV cannot be present in any control sample, CNVs are iteratively challenged with the Multiscale CNV detection algorithm against each control.

### Generation of simulated data

Heterozygous deletions and duplications in randomly picked exonic regions have been inserted in samples BAM files using the library Pysam from Python. Briefly, a script selects a random exonic position (inter-exonic or across two or multiple exons) and, around that location, removes or duplicates half of the overlapping reads in the sample BAM files, respectively. Then coverage (COV) files are produced with samtools depth following the same protocol as for the normal samples and processed with CoM with standard parameters.

## Data availability

CoverageMaster is available at https://github.com/fredsanto/coverageMaster.

## Acknowledgments

We thank Marco Belfiore from Genesupport for providing the array CGH data and Xavier Blanc from Medigenome for constructive discussions and suggestions. The study was partially supported by the Swiss National Science Foundation (310030_185292), Horizon2020 (miniNO 847941) and Novartis Foundation (18A052) to F.S.

## Bibliography

1. Shlien, A. & Malkin, D. Copy number variations and cancer. Genome Med 1, 62 (2009).

2. Truty, R. et al. Prevalence and properties of intragenic copy-number variation in Mendelian disease genes. Genet Med 21, 114–123 (2019).

3. Zack, T.I. et al. Pan-cancer patterns of somatic copy number alteration. Nat Genet 45, 1134–40 (2013).

4. Cancer Genome Atlas Research, N. et al. The Cancer Genome Atlas Pan-Cancer analysis project. Nat Genet 45, 1113–20 (2013).

5. Stuppia, L., Antonucci, I., Palka, G. & Gatta, V. Use of the MLPA assay in the molecular diagnosis of gene copy number alterations in human genetic diseases. Int J Mol Sci 13, 3245–76 (2012).

6. Rieber, N. et al. Coverage bias and sensitivity of variant calling for four whole-genome sequencing technologies. PLoS One 8, e66621 (2013).

7. Marshall, C.R. et al. The Medical Genome Initiative: moving whole-genome sequencing for rare disease diagnosis to the clinic. Genome Med 12, 48 (2020).

8. Sathirapongsasuti, J.F. et al. Exome sequencing-based copy-number variation and loss of heterozygosity detection: ExomeCNV. Bioinformatics 27, 2648–54 (2011).

9. Krumm, N. et al. Copy number variation detection and genotyping from exome sequence data. Genome Res 22, 1525–32 (2012).

10. Plagnol, V. et al. A robust model for read count data in exome sequencing experiments and implications for copy number variant calling. Bioinformatics 28, 2747–54 (2012).

11. Goodwin, S., McPherson, J.D. & McCombie, W.R. Coming of age: ten years of nextgeneration sequencing technologies. Nat Rev Genet 17, 333–51 (2016).

12. Parihar, R. & Ganesh, S. The SCN1A gene variants and epileptic encephalopathies. J Hum Genet 58, 573–80 (2013).

13. Ogino, S. & Wilson, R.B. Genetic testing and risk assessment for spinal muscular atrophy (SMA). Hum Genet 111, 477–500 (2002).

14. Van der Auwera, G.A. et al. From FastQ data to high confidence variant calls: the Genome Analysis Toolkit best practices pipeline. Curr Protoc Bioinformatics 43, 11 10 1–11 10 33 (2013).

15. Li, H. & Durbin, R. Fast and accurate short read alignment with Burrows-Wheeler transform. Bioinformatics 25, 1754–60 (2009).

16. Zarrei, M., MacDonald, J.R., Merico, D. & Scherer, S.W. A copy number variation map of the human genome. Nat Rev Genet 16, 172–83 (2015).

